# A quantitative model for characterizing the evolutionary history of mammalian gene expression

**DOI:** 10.1101/229096

**Authors:** Jenny Chen, Ross Swofford, Jeremy Johnson, Beryl B. Cummings, Noga Rogel, Kerstin Lindblad-Toh, Wilfried Haerty, Federica di Palma, Aviv Regev

## Abstract

Characterizing the evolutionary history of a gene’s expression profile is a critical component for understanding the relationship between genotype, expression, and phenotype. However, it is not well-established how best to distinguish the different evolutionary forces acting on gene expression. Here, we use RNA-seq across 7 tissues from 17 mammalian species to show that expression evolution across mammals is accurately modeled by the Ornstein-Uhlenbeck (OU) process. This stochastic process models expression trajectories across time as Gaussian distributions whose variance is parameterized by the rate of genetic drift and strength of stabilizing selection. We use these mathematical properties to identify expression pathways under neutral, stabilizing, and directional selection, and quantify the extent of selective pressure on a gene’s expression. We further detect deleterious expression levels outside expected evolutionary distributions in expression data from individual patients. Our work provides a statistical framework for interpreting expression data across species and in disease.

**One Sentence Summary:** We demonstrate the power of a stochastic model for quantifying selective pressure on expression and estimating evolutionary distributions of optimal gene expression.

---

Comparative genomics has identified and annotated functional genetic elements by their evolutionary patterns across species (*1–6*). Current comparative studies focus primarily on analysis of genomic sequences, using a well-established theoretical framework developed in the 1960s based on observations that neutral sequence diverges linearly across time (*7–11*). These models allow for detection of sequence elements that evolve slower (*e.g.*, due to purifying selection) or faster (*e.g.*, due to positive selection or relaxed selective constraints) than expected under the null model of neutral evolution.

It has long been accepted that divergence of gene regulation, manifested by phenotypic changes in gene expression, also plays a key role in evolution (*12–17*). An evolutionary analysis of gene expression should help interpret gene function and evolutionary processes in ways that cannot be addressed by sequence alone: signatures of purifying selection on a gene’s expression across tissues could reveal the context in which it plays the most important role; the strength of evolutionary constraint on a gene’s expression could help interpret expression data from clinical samples; and genes whose expression is under directional (positive) selection can help assess the basis of lineage- and species-specific phenotypes.

Though theoretical models of expression evolution have been proposed (*18–21*), most studies have not applied these models to address functional questions. Initial studies on an evolutionary model for gene expression have also been hampered by small datasets, leading to conflicting results. Indeed, some studies concluded that expression is diverging at a linear rate across time, suggestive of widespread neutral evolution (*22–25*), whereas others determined that the evolutionary rate is constant, suggestive of strong purifying selection (*20, 26–29*).

To systematically explore expression evolution, we compiled a well-sampled dataset across the mammalian phylogeny, spanning 17 species and 7 different tissues (brain, heart, muscle, lung, kidney, liver, testis) (**Fig. 1A, table S1**). The dataset combines published data for 12 species (*28, 30–34*) with data for five additional species we newly collected here (**Fig. 1A, asterisk, SOM**) to improve phylogenetic coverage. We focused on the 10,899 annotated mammalian one-to-one orthologs (*35*). As expected (*36*), expression profiles first cluster by tissue and then by species (fig. S1), and their hierarchical clustering closely matches the phylogenetic tree (**fig. S2**).

**Figure 1.**
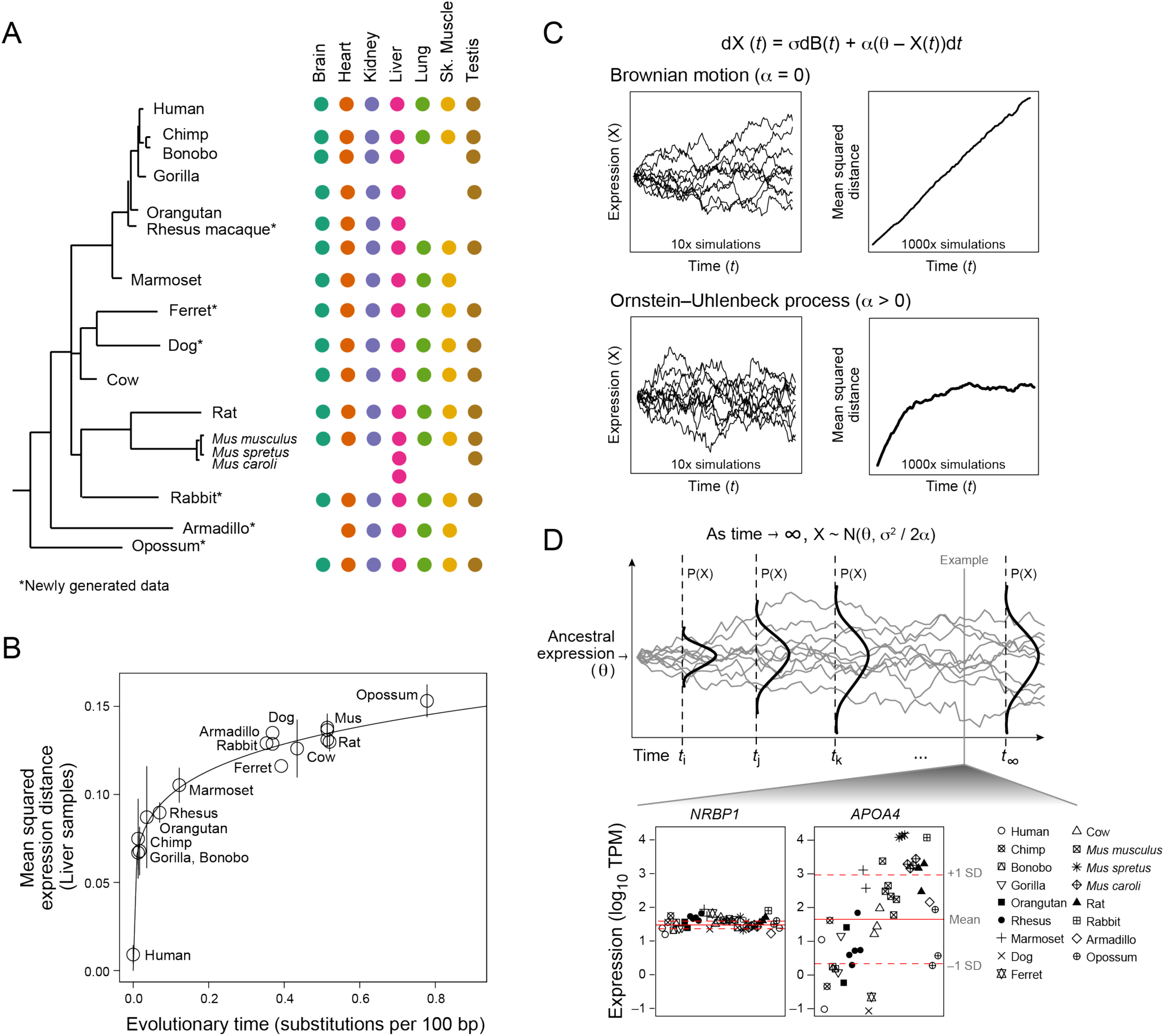
Expression evolution across mammalian lineages is accurately modelled by the OU process. (**A**) Data overview. Phylogenetic tree of all 17 mammals (left) marked by tissue types (colored dots, right) for which profiles are included. Asterisk: Newly generated data. (**B**) Expression divergence is not linear. Shown is the pairwise mean squared expression distances (y-axis) between mammals and human for liver samples across evolutionary time, as estimated by substitutions per 100 bp (x-axis). Error bars: standard deviation of the mean across replicates. Solid line: Nonlinear regression fit. (**C**) OU model. Top: Equation describing OU model (σ: rate of genetic drift; dB(t): Brownian motion; θ: optimal expression level; α: strength of selection). Left: Simulated trajectories of expression (y-axis) over evolutionary time (x-axis) under a Brownian motion (top) and OU (bottom) process. Ten example trajectories are shown. Right: Mean squared distance to initial value (y-axis) across time (x-axis) from 100 simulated trajectories. (**D**) Distribution of optimal expression. Top: Illustration of the change in probability distribution of expression (y-axis) across time (x-axis) under an OU process. The distribution stabilizes as time approaches infinity. Bottom: Scatter plot of log_10_(TPM) values (y-axis) across all liver samples (x-axis) of two example genes with low (*NRBP1*) and high (*APOA4*) variance. Solid and dotted red lines: estimated mean and variance, respectively, of the asymptotic (optimal) distribution of each gene’s expression value estimated using the OU process. Note that mean and variance are calculated in log space.

On average, pairwise expression differences relative to a specific species (SOM) do not increase linearly with evolutionary time (**Fig. 1B, fig. S3,4**), but instead plateau. For example, when comparing to human profiles, differences plateau beyond the primate lineage (43.2 million years) (*37*) (**Fig. 1B, fig. S3**). This relationship is observed in each of the five tissues for which we have expression data for all primates (brain, heart, kidney, liver, testis) (**fig. S3**), and regardless of reference species (**fig. S4**), albeit with varying rates.

The observed pattern of expression divergence is accurately modelled by an Ornstein-Uhlenbeck (OU) process (average mean squared error across tissues = 0.002) (**Fig. 1C, D**). This stochastic process was initially proposed as a model for phenotypic evolution by Hansen (*19*) and has more recently gained traction for modelling expression evolution across *Drosophila*, suggesting that it may be generalizable across the animal kingdom (*20, 21, 38*). Thus far, OU models have primarily been employed for theoretical inferences about fitness gains and selective effects of evolving expression levels, with one previous limited application for detecting selection on expression across a smaller mammalian phylogeny of 9 species (*28*). Here, we extend previous work with a better powered phylogeny to demonstrate how to apply the OU model to yield interpretable answers to functional questions about expression evolution, disease gene discovery, and lineage-specific adaptations.

In the context of expression, the OU process (**Fig. 1C**) is a modification of a random walk, describing the change in expression (dX_t_) across time (dt) by dX_t_ = σdB_t_ + α(θ - X_t_) dt, where dB_t_ denotes a Brownian motion process. The model elegantly quantifies the contribution of both drift and selective pressure for any given gene: (**1**) drift is modeled by Brownian motion with a rate σ (**Fig. 1C**, top), while (**2**) the strength of selective pressure driving expression back to an optimal expression level θ is parameterized by α (**Fig. 1C**, bottom). The OU process incorporates time information and fully accounts for phylogenetic relationships, thus allowing us to fit individual evolutionary expression trajectories. At longer time scales, the interplay between the rate of drift (σ) and the strength of selection (α) reaches equilibrium and, as time increases to infinity, constrains expression X_t_ to a stable, normal distribution, with a mean, θ, and variance, σ^2^ / 2α (**Fig. 1D**).

We applied the OU model to yield biologically interpretable results to evolutionary questions about gene expression. **First**, for each tissue separately, we estimate from our data, the asymptotic distribution of optimal expression for genes under stabilizing selection. We then use this distribution’s OU variance (which we term ‘evolutionary variance’) to characterize how constrained a gene’s expression is in each tissue. **Second**, we compare the observed gene expression in an individual patient with disease to the optimal level from the model, in order to detect potentially deleterious expression levels and use those to nominate causal disease genes. **Third**, we use an extension of the OU model (*39*) to account for the existence of multiple distributions of optimal expression within a phylogeny and apply them to identify genes undergoing adaptive evolution within subclades. We describe each of these applications in turn.

To test whether a gene is under stabilizing selection, we used a likelihood ratio test to compare the fit with no selection (α = 0; Brownian motion only; **Fig. 2A**, top) to one with stabilizing selection (α > 0, OU process; **Fig. 2A**, bottom). On average, 83% of genes (range: 78% - 87%; FDR < 0.05) were under stabilizing selection (**Fig. 2A, bottom, fig. S5**). Nevertheless, the expression of hundreds of genes within each tissue appeared to be neutrally evolving (Fig. 2A, top, fig. S5). Comparing across tissues and considering only genes robustly expressed over 5 transcripts per million (TPM), 57% (5,669/8,913) of all genes were under stabilizing selection in *all* tissues in which they were expressed, 39% (2,722) were under stabilizing selection in only some of the tissues where they were expressed, and only 6% (521) were not under stabilizing selection in any of the tissues in our study (**Fig. 2B**).

**Figure 2.**
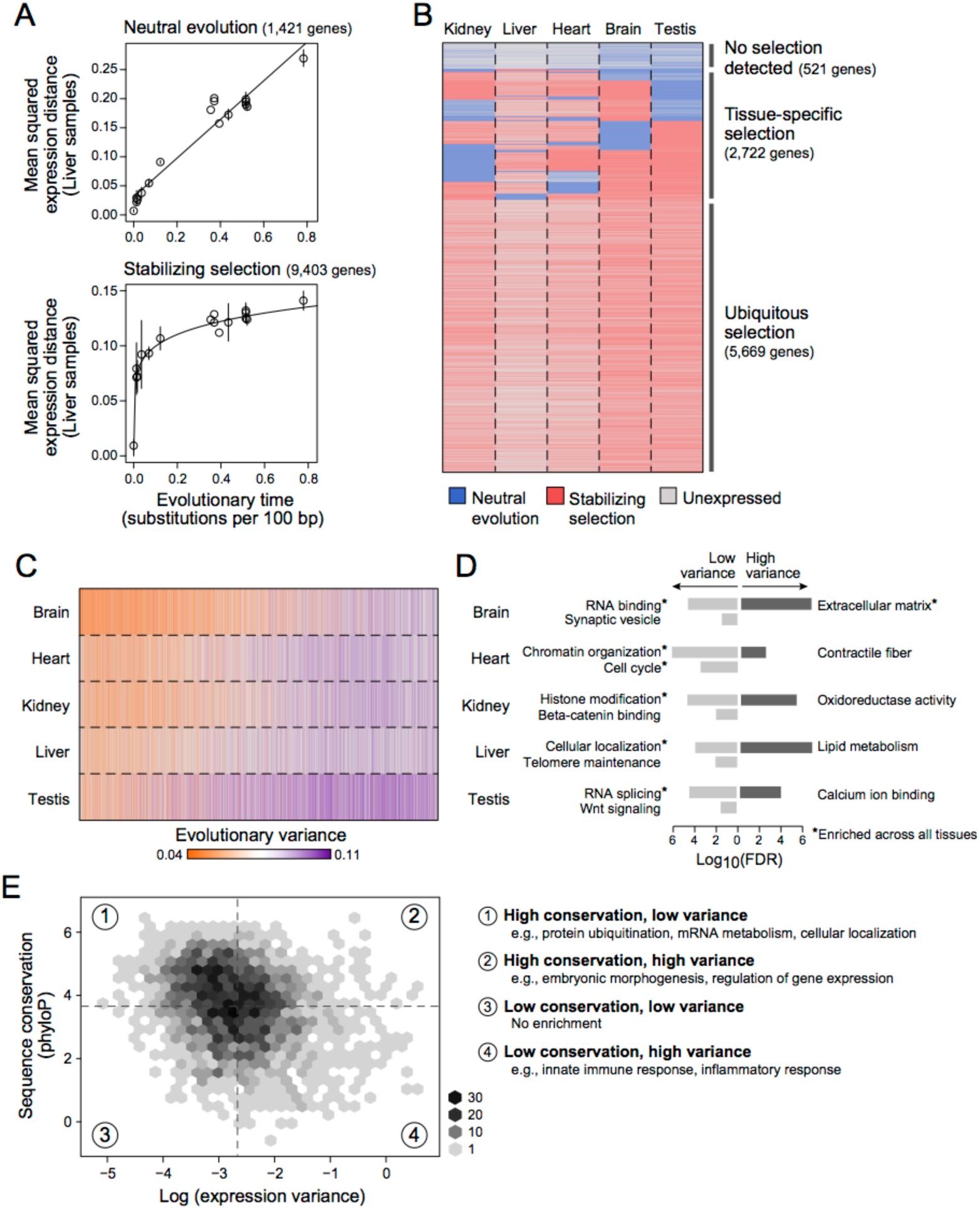
Quantification of neutral and constrained selection on gene expression using the OU model parameters. **(A)** Detection of stabilizing selection. Pairwise mean squared expression distances (y-axis) between mammals and human for liver samples across evolutionary time (x-axis) for 1,421 genes whose expression evolution fits better under a Brownian motion (BM) process (top), indicating neutral evolution, and 9,403 genes whose expression evolution fits better a Ornstein-Uhlenbeck (OU) process (bottom), indicating the presence of stabilizing selection. Solids lines: Linear regression fit for BM genes and nonlinear regression fit for OU genes. **(B)** Neutral and stabilizing selection across genes and tissues. Heatmap indicating genes (rows) whose expression is predicted to be evolving under neutral evolution (blue) or stabilizing selection (red) across 5 different tissues (columns). Gray: Genes that are expressed < 5 TPM. (C, D) Evolutionary variance across tissues and processes. (**C**) Heatmap shows estimated evolutionary variance of expression (colorbar: orange: low; purple: high) across 8,794 genes (columns) in 5 tissues (rows). Gray: Genes expressed < 5 TPM. (**D**) Barplot of −log_10_FDR values for significantly enriched GO categories of low (light gray) and high (dark gray) variance genes within each tissue. Asterisk: Category enriched in every tissue. (**E**) Relation between sequence and expression evolution. Binned scatterplot of log(evolutionary variance) of liver expression (x-axis) vs. sequence conservation, as measured by the phyloP score (y-axis). Median variance and phyloP scores are indicated by vertical and horizontal dotted lines, respectively. Enriched GO categories (FDR < .001) for genes in each quadrant of the scatterplot are listed on the right.

We assessed the sensitivity and specificity of our detection of genes under expression-stabilizing selection using a jackknifing procedure, where we subsampled to consider phylogenies ranging from 3 to 16 species (**SOM**). As expected, the number of genes called under stabilizing selection *(i.e.,* rejecting the null hypothesis) increases as more species are included (**fig. S6A**), but does saturate at 14 species. Importantly, the false positive rate (relative to analysis of the full dataset) is very low: less than 1% of genes found as under selection with a subsampled phylogeny are found to be neutral (i.e., accepting the null hypothesis) with the full phylogeny (**fig. S6B**). The OU model’s ‘evolutionary variance’ is highly robust to subsampling, as determined by the very low mean squared error (MSE < 0.005) when estimating variance from subsampled phylogenies. Indeed, with less than 6 species, the evolutionary variance is far more robust than the simple sample variance used by non-phylogenetic methods (**fig. S6C**). Note that lowly expressed genes (< 5 TPM) had very high evolutionary variance (**fig. S7**), likely due to technical noise (*40*), and were thus excluded from all subsequent analyses, resulting an median of 5,860 analyzed genes per tissue.

The OU model’s evolutionary variance quantifies the extent of evolutionary constraint on a gene’s expression in each tissue, reflecting distinctions between tissues, genes and processes. Brain had the most genes with low variance (most constraint), and testis the least, consistent with previous estimates of rate of expression evolution for those tissues (*28, 41*) (**Fig. 2C and fig. S8**). Variance was highly correlated between somatic tissues (mean Pearson’s *r* = 0.84), and less correlated between somatic tissues and testis (mean Pearson’s *r* = 0.55) (**fig. S9A**). For genes expressed across three or more somatic tissues, expression level across tissues was negatively correlated with variance across the tissues (median Pearson’s *r* = −0.27), though the tissue of highest expression only matched tissue of lowest variance in 34.5% (1,673 / 4,840) of genes (**fig. S9B**).

Evolutionary variance and function were strongly associated, consistent with previous reports (*42*): across all tissues, genes with low variance were enriched for housekeeping functions (*e.g.*, RNA binding and splicing, chromatin organization, cell cycle), whereas those with high variance were enriched for extracellular proteins (rank based enrichment test, **SOM**, FDR < 0.001). Some processes were enriched in genes with low or high variance only in specific tissues (**Fig. 2D, table S2**): among the conserved (low variance) processes were synaptic proteins in brain (FDR = 0. 011) and Wnt signaling in testis (FDR = 0.014); processes with high variance included contractile fiber part in heart (FDR = 0.005), oxidoreductase activity in kidney (FDR = 6.10*10^−6^), and lipid metabolism in liver (2.31*10^−9^).

There was only a modest correlation between expression and sequence conservation (Pearson’s *r* = −0.25) (**Fig. 2E, SOM**). Genes conserved in both expression and sequence were significantly enriched for housekeeping processes (FDR < 10^−4^**, Fig. 2E, table S3**), and genes divergent in both were enriched for immune and inflammatory response (FDR < 10^−6^, **Fig. 2E, table S3**). More intriguingly, genes conserved in sequence but divergent in expression were enriched in transcriptional regulators (FDR = 3.1*10^−5^), especially those involved in embryonic morphogenesis (FDR = 9.8*10^−8^; *e.g., IRX5, HAND2, NOTCH1*). Although higher variance may be impacted by environment, changes in cell type composition, and genetic differences, our analysis supports the hypothesis that divergence in gene regulation without protein sequence divergence can account for species-specific phenotypes.

In analysis of rare diseases, sequence conservation is commonly used to prioritize mutations in genes that are more essential and likely causal for rare diseases when mutated (*43–45*). By analogy, we hypothesized that expression conservation should also be predictive of gene essentiality. Indeed, genes that are either essential in culture (*46*), essential in mice (*47*), or haploinsufficient in humans (*48*) had significantly lower evolutionary variance (higher constraint) than their non-essential or haplosufficient counterparts across almost all tissues (Wilcoxon rank-sum test p-value < 0.01, **Fig. 3A**). Moreover, disease genes with tissue-specific expression (> 5 TPM in three or fewer tissues) consistently exhibited significantly lower variance in the disease-relevant tissue than tissue-specific non-disease genes in each of three tested settings: rare single genes directly linked to non-syndromic autism spectrum disorder (ASD) (brain) (*49*), congenital heart defects (heart) (*50, 51*), and neuromuscular disease (skeletal muscle) (*52*) (p-value < 0.05, **Fig. 3B**). In ASD-linked genes (but not the other two conditions), we also observed significantly lower variance of ubiquitously expressed disease vs. non-disease genes, perhaps related to observed high rates of co-morbidities with ASD (*53*).

**Figure 3.**
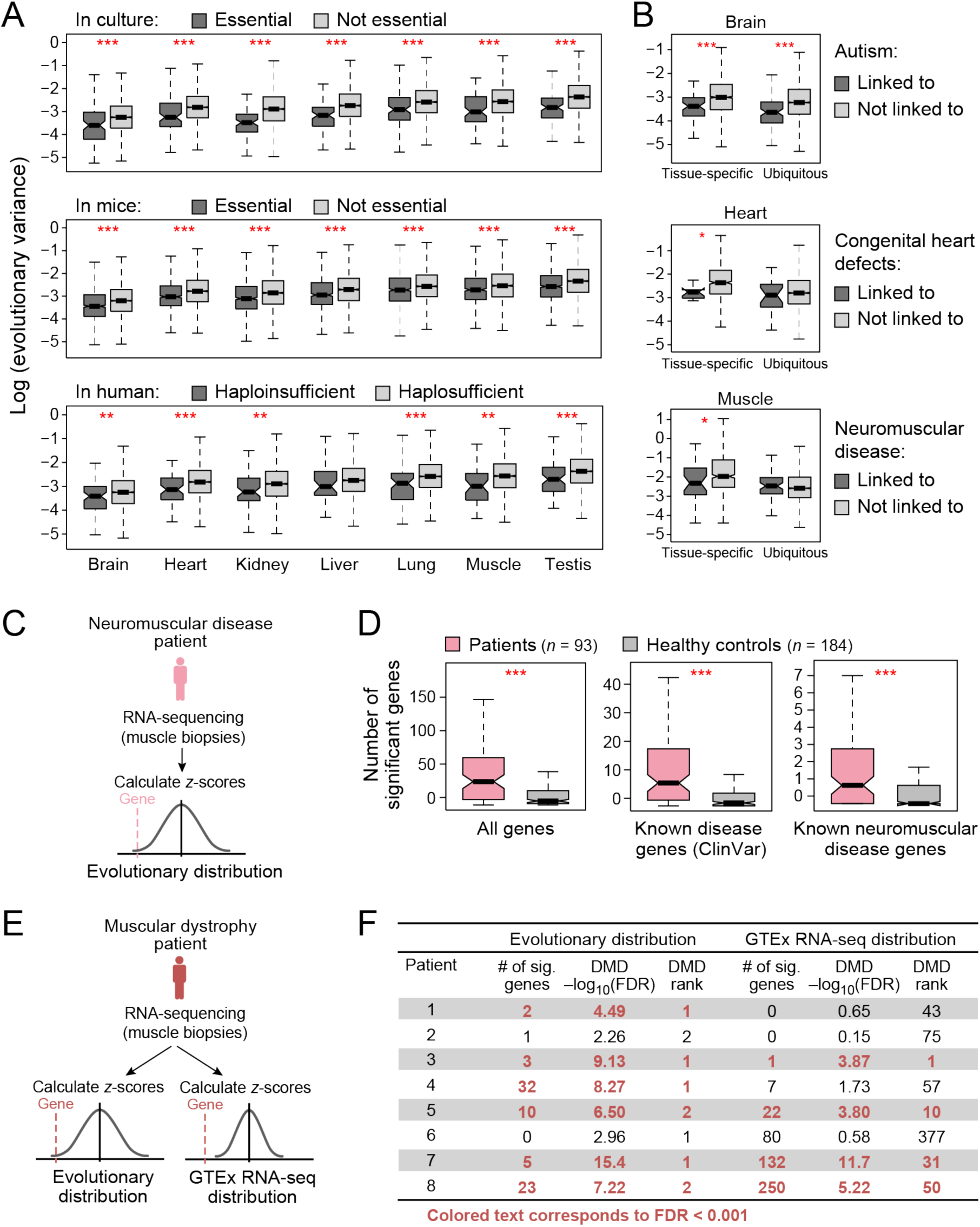
Evolutionary distribution of gene expression helps identify disease-contributing genes. **(A)** Essential genes have lower evolutionary variance. Boxplots show the distribution of log(evolutionary variance) (y-axis) of genes essential in culture (top), essential in mice (middle), and haploinsufficient in human (bottom) (dark gray), and their non-essential or haplosufficient counterparts (light gray) in each of 7 tissues (x-axis). *** denotes *p* < 0.001 and ** denotes *p* < 0.01. (**B**) Disease genes have lower evolutionary variance. Boxplots show the distribution of log(evolutionary variance) (y-axis) of genes linked (dark gray) and not linked (light gray) to high-penetrance monogenic autism spectrum disorder (top), congenital heart defects (middle), and neuromuscular disease (bottom) in the relevant tissue (brain, heart, and muscle, respectively). Left boxes: genes that are specifically expressed (> 5 TPM in 3 or fewer tissues) in that tissue; right boxes: genes that are ubiquitously expressed. (**C**) Overview of using evolutionary distributions to identify outlier gene expression in RNA-seq of patient muscle biopsies. *** denotes *p* < 0.001 and * denotes *p* < 0.05. (**D**) Outlier expression in disease tissue. Boxplots of the number of genes with significant z-scores (FDR < 0.01) from neuromuscular patient data (pink) and from healthy controls (gray) when considering all genes (left), known disease genes from ClinVar (*58*) (middle), and known neuromuscular disease genes (right). *** denotes *p* < 0.001. (**E, F**) Using evolutionary distributions or GTEx RNA-seq distributions to identify outlier gene expression from RNA-seq of muscular dystrophy patients. (**E**) Two scoring approaches based on evolutionary distributions (left) or GTEx RNA-seq distributions (right). (**F**) Table shows number of significant outlier genes, −log_10_(FDR) score and *DMD*’s rank for outlier gene expression for all patients with muscular dystrophy when using distributions estimated from evolutionary data (left) or GTEx RNA-seq (right).

Next, we hypothesized that the parameters of the OU model can predict disease genes, by highlighting outlier, likely pathogenic, gene expression patterns in rare disease patient data by comparing patient expression to each gene’s optimal OU distributions (**Fig. 3C**). This is analogous to causal disease gene discovery by identifying putatively pathogenic sequence mutations in whole exome sequencing (*54–57*). To this end, we obtained RNA-seq of muscle biopsies of 93 patients clinically diagnosed with neuromuscular disease (**SOM, table S4**). For each patient sample, we calculated a z-score for each gene to assess how they deviate from the (optimal) evolutionary fit for that gene’s expression in skeletal muscle, with correction for multiple hypothesis testing (**Fig. 3C, SOM**). Compared to GTEx muscle samples from 184 healthy people (*58*), patients had, on average, 3.2-fold more dysregulated genes overall by this measure (Wilcoxon rank sum test *p*-value =2*10^−9^, **Fig. 3D**, left), 3.0-fold more dysregulated muscle-expressed disease genes (*59*) (*p*-value = 2.1*10^−10^, **Fig. 3D**, middle), and 2.0-fold more dysregulated known neuromuscular disease genes (*p*-value = 2.7*10^−4^, **Fig. 3D**, right). This suggests that the evolutionary parameters fit by the OU model can be used to detect outlier expression values that are more likely to be deleterious. Importantly, in contrast to methods for differential expression between patient and healthy controls, the test does not require a control population, and can be conducted for a single patient sample.

We then tested whether the OU model could be used to identify the causative gene in rare disease analysis. As a proof of principle, we focused on the subset of 8 patients from the muscle disease cohort who were clinically diagnosed with either Becker or Duchenne muscular dystrophy, including confirmation of absent or decreased dystrophin protein via immunoblotting (*52*). To compare our approach to a standard differential expression analysis, we ranked genes by outlier expression with z-scores defined based either on (**1**) comparison to the mean and variance estimated from our evolutionary data; or (2) comparison to a mean and variance estimated from only healthy GTEx human data (**Fig. 3E**). By our evolutionary data, fewer genes ranked as significant outliers in each patient (median: 4, range: 0 – 32), and the *DMD* gene ranked as either the top or second most significantly aberrantly expressed gene in 6 of 8 patients, each showing significant underexpression (FDR < 10^−3^) (**Fig. 3F**). By comparison, scoring in reference to GTEx expression data did not yield such specific results: a median of 14.5 genes were outliers (range: 0 - 250), only 4 of 8 patients were called as significantly underexpressing *DMD* (FDR < 10^−3^), and its significance in these patients ranked between 1 and 50. Thus, using the OU model’s estimate of evolutionary mean and variance of optimal gene expression helps detect gene dysregulation of the actual disease gene and could aid novel disease gene discovery in individual patients, even without any control samples.

Finally, we applied an extension of the OU model (*39*) to detect directional selection in gene expression. To enrich for changes that are likely to have resulted from ancestral genetic changes, rather than environmental causes, we focused on detecting shifts in expression consistent in direction and magnitude across entire subclades of two or more mammals. We used an extension of the OU model with multiple selection regimes across a single phylogeny (*39*), by determining the distribution of expression level as a *multivariate* normal distribution whose mean and variance are estimated for each (predefined) subclade (**Fig. 4A**). We then identified “differential gene expression” across the tree, in the following way: We applied the extended model for each gene in each tissue and tested each of four alternative hypotheses: OU_all_, which models a single optimum for all species, and OU_primates_, OU_rodents_, OU_carnivores_, each modeling two optima, one for the ancestral distribution and one for the distribution within primates (branch length = 0.12), rodents (branch length = 0.18), or carnivores (branch length = 0.20), respectively (**Fig. 4B**). For each gene, we first used a likelihood ratio test between each OU model and the null hypothesis of a Brownian motion model and removed any models against which the null model could not be rejected. We then assigned the best OU model using goodness-of-fit tests (only when both Akaike and a Bayesian information criterion scores agreed on the best fitting model, **SOM**). Finally, as a conservative measure, we retained only those genes that also changed at least 2-fold between subclades and had a mean expression level of at least 5 TPM in one of the subclades.

**Figure 4.**
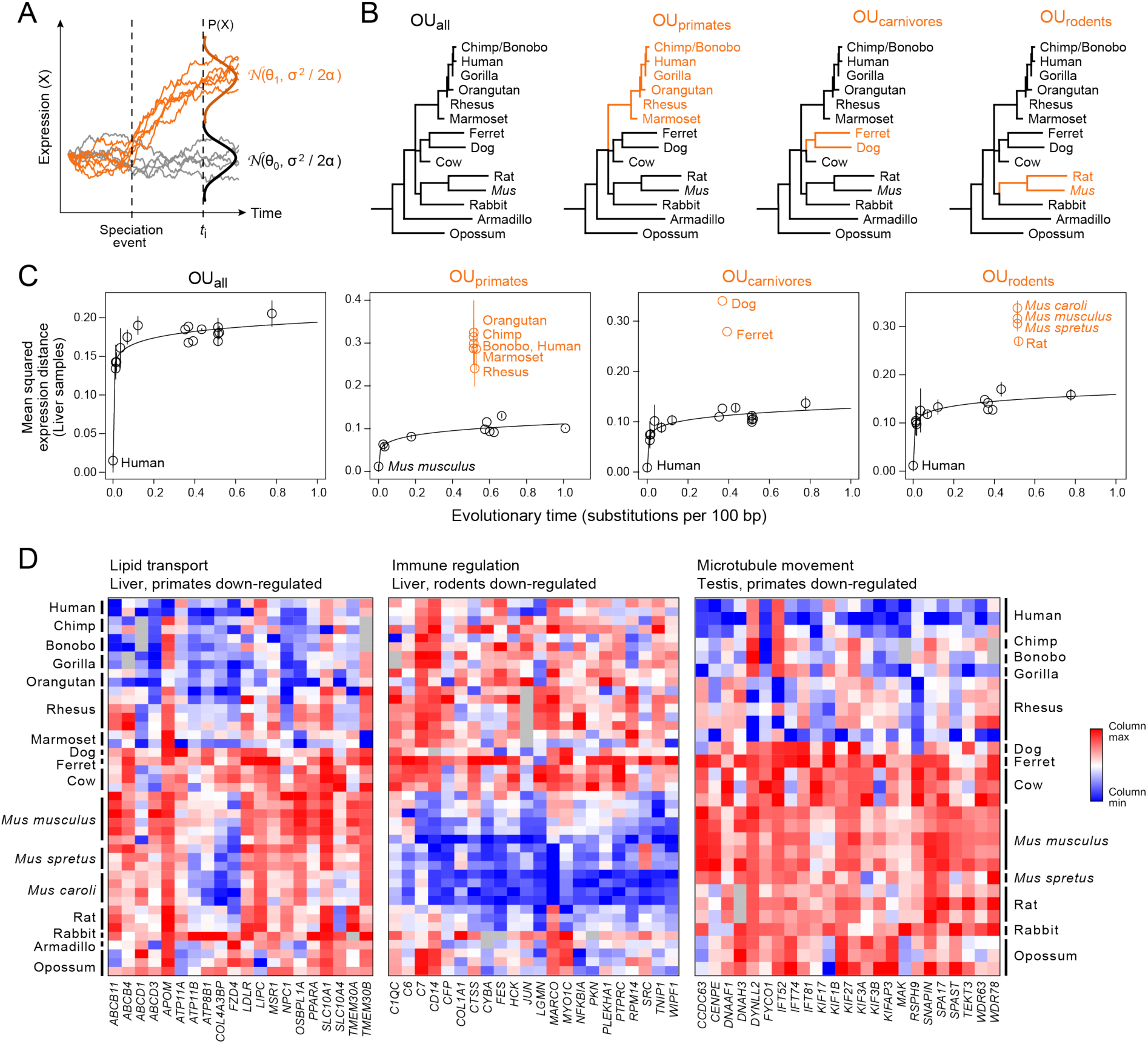
Multivariate OU process enables detection of lineage-specific expression changes. (**A**) Multivariate OU process. Simulated trajectories of expression (y-axis) over time (x-axis) under a multivariate Ornstein-Uhlenbeck (OU) process. Trajectories in gray are sampled from the same distribution (𝓝_0_) across time while trajectories in orange start at the same ancestral distribution (𝓝_0_) but evolve under a new distribution (𝓝_1_) after a speciation event. (**B**) Four tested hypotheses of expression evolution. From left: the univariate OU_all_ model, in which gene expression evolves under a single stabilizing regime across the phylogeny (black), and three multivariate OU models, OU_primates_, OU_carnivores_, and OU_rodents_, in which gene expression evolves under the ancestral regime (black) and a new regime in the specified subclade (orange). (**C**) Lineage specific expression in liver. Pairwise mean squared expression distances (y-axis) between a reference species (labeled black point) and each of the other mammals in liver samples for genes assigned to each of four tested OU models. Black points: Species evolving under ancestral distribution; Labeled orange points: species evolving under new regime after the lineage split. Solid line: nonlinear regression fit for species evolving under ancestral distribution. (**D**) Example processes enriched for lineage-specific expression. Heatmaps show column-normalized expression (colorbar) from genes (columns) with lineage-specific expression patterns in specific tissues in three enriched gene ontology categories (FDR < 0.05): lipid transport in liver (left), immune regulation in liver (middle), and microtubule movement in testis (right).

Within each somatic tissue, we found about 26% of expressed genes to be lineage-specific, while strikingly, in testis, we found almost half (48%) of expressed genes appeared to have lineage-specific differential expression. However, when we used a control analysis with shuffled species assignments to estimate false discovery rates (**SOM**), we found that we achieved FDR < 30% in only liver (all three clades) and testis (two of three), as well as in the primate clade for brain and lung (**fig. S10**). As an example, in liver, we identified 640, 794, and 615 genes with lineage-specific expression changes in primates, rodents, and carnivores, respectively, highlighting specific metabolic processes diverging in regulation in each clade. The lineage-specific genes deviated significantly from expectation only if there was no clade-specific selection (**Fig. 4C**). Testing for functional enrichment among lineage-specific genes (**table S5**), we found primate-specific downregulation of genes related to a number of lipid metabolic processes in the liver (FDR = 1.88*10^−11^). These processes include peroxisomal functions (FDR = 2.45*10^−8^), fatty acid metabolism (FDR = 1.52*10^−8^), and lipid transport (FDR = 3.36*10^−3^) (**Fig. 4D**, table S5), and include known regulators of lipid metabolism such as the LDL receptor (*LDLR*) (*60*), hepatic lipase (*LIPC*) (*61*), and the transcription factor *PPAR-alpha* (*62*). Thus, the expression of multiple pathways may have diverged at the ancestral primate branch, consistent with observations that human lipidemia is not well-modeled by mice without further genetic modification (*63*). In another example, genes involved in regulation of immune response were downregulated across rodent livers (FDR = 6.97*10^−4^), and in testis, microtubule-based movement genes (FDR = 2.82*10^−3^) and spermatogenesis (FDR = 2.82*10^−2^) were downregulated across primates (**Fig. 4D**), reflecting the known rapid evolution of immune-(*42, 64, 65*) and reproduction-related genes (*66, 67*).

In conclusion, by combining a large dataset of comparative gene expression profiles across mammals with systematic analysis, we showed that gene expression of one-to-one mammalian orthologs is evolving nonlinearly across evolutionary time and is accurately modeled by an OU process. We then show how to use this model to answer three key questions: (**1**) testing individual gene level for selective regimes, including purifying selection and extent of constraint, (**2**) identifying deleterious gene expression in single patient disease tissue by characterizing outliers relative to a predicted *distribution* of optimal expression for each gene, and (**3**) detecting lineage-specific expression using an extension that accounts for multiple distributions of optimal expression. Looking forward, we anticipate that the OU model can be further developed for other biological queries, for example by incorporating the contribution of within-species expression variation (*68*) or testing for stabilizing selection across pathways of genes or paralog families. As shown by our analysis, characterizations of expression across additional tissue types and species under varied developmental and environmental contexts will provide increased power and further insight into the evolution of gene expression, and the relationship between genotype and phenotype.

## Materials and Methods

### Data collection

The following table summarizes the sources for all data used in this study. For a more detailed table of SRA accession numbers and read alignment statistics, see **Table S1**.

#### Sources

Brawand et al. 2011 (*28*)

Illumina Body Map 2.0 at https://www.ebi.ac.uk/arrayexpress/experiments/E-MTAB-513/ Non-human primate reference transcriptome resource (NHPRTR) (*69*)

Merkin et al. 2012 (*30*)

Cortez et al. 2014 (*32*)

Wong et al. 2015 (*33*)

Harr and Turner 2010 (*34*)

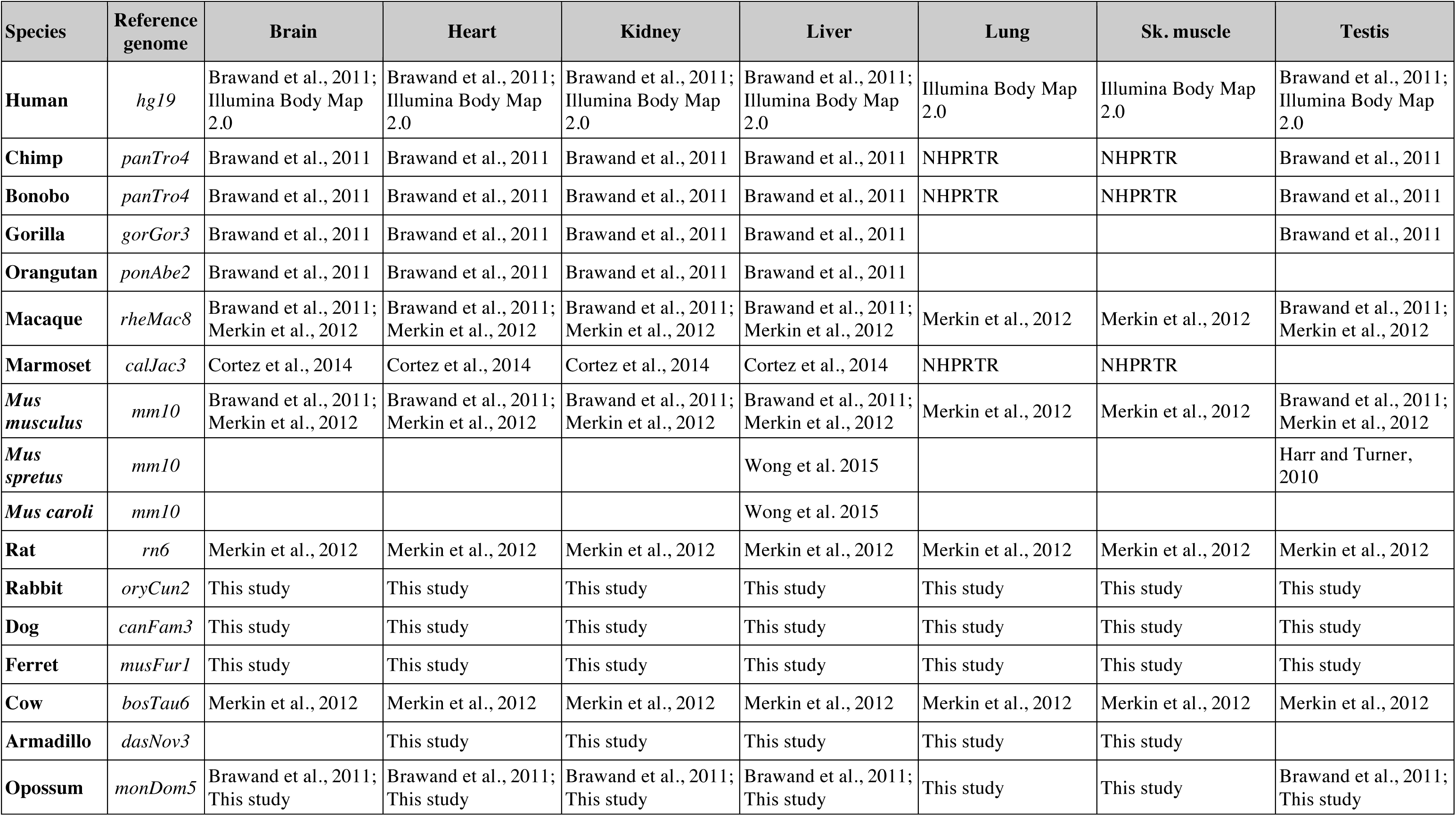

### Samples for evolutionary dataset

RNA samples from dog and rabbit tissues were commercially obtained from Zyagen. RNA samples from opossum tissues were a kind gift from Paul Samallow (Texas A&M). RNA samples from armadillo tissues were a kind gift from Jason Merkin and Christopher Burge (MIT). All tissue collection was approved by IACUC and carried out in accordance with respective institutional guidelines.

### RNA-sequencing for evolutionary dataset

RNA-sequencing (RNA-seq) libraries were prepared as described in (*70*). Briefly, 10 μg total RNA was poly-A selected twice using Dynabeads mRNA Purification Kit (Invitrogen, 610.06). Resulting mRNA was DNase treated (Ambion AM1907) and then fragmented using heat. First strand cDNA synthesis was performed using the SuperScript Double-Stranded cDNA Synthesis Kit (Invitrogen, 11917-010), supplementing in SuperScript III Reverse Transcriptase (Invitrogen, 18080-093), incorporating SUPERase*In (Ambion, AM2694), and Actinomycin D (USB, 10415). First strand cDNA was cleaned using 1.8X RNAClean XP SPRI beads (Beckman Coulter, A64987). Second-Strand Synthesis was performed replacing dTTP with dUTP, and the resulting double-stranded cDNA was cleaned using a MinElute PCR Purification Kit (Qiagen, 28004). Illumina libraries were constructed by repairing the ends of the cDNA, ligating adapters, and cleaning/size-selecting with 0.7x SPRI. Illumina libraries were treated with USER to excise dUTP, and amplified via PCR using Fusion Master mix with GC buffer (NEB, F532S). Samples were sequenced on an Illumina HiSeq 2000 sequencer, to a minimum depth of 35 million reads.

### Genome and transcriptome annotations

All genomes were downloaded from the UCSC Genome Browser (*71*). To assembl transcriptomes, Ensembl gene annotations (*35*) were downloaded from UCSC Table Browser (table **ensGene**) and converted to sequence using BEDTools (*72*). Ortholog annotations were downloaded from Ensembl BioMart (Ensembl Genes 90) (73). Only genes that met the following criteria were used for this study: (**1**) no duplications in any of the studied mammals, (**2**) an ortholog present in either armadillo or opossum (*i.e.* placental mammal or marsupial outgroup), (**3**) no more than 3 deletions across primates (human, chimp, gorilla, orangutan, macaque, marmoset), (**4**) no more than 1 deletion across glires (mouse, rat, rabbit), and (**5**) no more than 1 deletion across laurasiatherians (cow, dog, ferret).

### Phylogenetic tree

The phylogenetic tree of vertebrate species was downloaded from UCSC Genome Browser at http://hgdownload.cse.ucsc.edu/goldenpath/hg19/multiz100way/ (*71*). Distances between mammals used in this study were extracted using the Environment for Tree Exploration Toolkit (*74*).

### Alignment and expression quantification

RSEM v1.2.12 (*75*) was used to align reads to the transcriptome of each species and to quantify transcripts per million (TPM) of each gene using default parameters.

### Modelling expression evolution

#### Quantifying expression difference

To calculate pairwise expression differences between each species (“comparing species”) and a reference species (*e.g.*, human in **Fig. 1**, or mouse in **fig. S3**), we applied principal component analysis (PCA) on pairwise gene expression levels (log_10_(TPM)), considering only genes that were expressed (> 0 TPM) in at least one species. For replicated samples (of the same tissue and species), we used the mean expression across replicates. For each tissue and each pair of species, we used the first principal component as the best fit line between the two species’ expression profiles. We then defined the pairwise expression difference as the orthogonal distance from the observed expression level in the comparing species to the best fit line. We used PCA rather than a linear regression because PCA accounts for noise in expression values from *both* species, while the linear regression would only model noise in the comparing species and treat the reference species as an independent variable.

#### Fitting linear and nonlinear regression models to mean squared expression distances

Under an Ornstein Uhlenbeck model, the expected mean squared distance across time follows a power law relationship (*y* = *ax^k^*). To fit this relationship between our observed mean squared expression distances (*y*) and evolutionary time (*x*), we log-transformed both axis to relate the variables linearly: log(*y*) = log(*a*) + *k*log(*x*). We then used basic least squares regression to find coefficients *a* and *k*.

For genes whose expression evolution fit better under a Brownian motion model (see below), we simpy used basic least squares regression to find the best fit line between mean squared expression distances and evolutionary time.

#### Fitting Ornstein-Uhlenbeck (OU) process parameters

Gene expression values (log_10_(TPM)) were first normalized within each tissue to human expression using TMM normalization (*76*) from the Bioconductor package *edgeR (77)*.

Brownian motion (BM) and Ornstein-Uhlenbeck (OU) models were fit to expression values using the R package *ouch* (*39*) with default parameters. P-values for each gene were calculated using a likelihood ratio test comparing the OU (alternative hypothesis) to the BM (null hypothesis) model, and then corrected for multiple hypothesis testing using the Benjamini-Hochberg false discovery rate (FDR) procedure (*78*).

#### Jackknifing procedure for estimating robustness of OU process parameters

To test the robustness of the OU model, we used a jackknifing procedure where we subsampled phylogenies ranging from 3 to 16 species (out of a total of 17 species). For each phylogeny size, we created ten randomly subsampled phylogenies. We then fit the OU model as described above.

### Measuring sequence conservation

Sequence conservation of a gene was defined by mean phyloP score (*3*) across the coding region of the longest annotated transcript of that gene.

### Gene ontology (GO) enrichment analysis

#### Relationship with evolutionary variance

To test for enriched GO categories across genes with low or high evolutionary variance, we us the ranked enrichment test from GOrilla (*79*). To avoid biases due to relationship between lowly expressed genes and high evolutionary variance estimates, we only used genes expressed at > 5 TPM.

#### Relationship with evolutionary variance and sequence conservation

We tested for enriched GO categories across genes in all four categories of high or low evolutionary variance and high or low sequence conservation. For each tissue separately, we defined “high” or “low” based on the median evolutionary expression variance and median phyloP score, respectively, and assigned all genes expressed at > 5 TPM to one of four categories. For GO enrichment analysis, where only sets with relatively large numbers of genes are typically enriched at levels that survive multiple hypothesis testing correction, we first unified the genes of each category across all tissues and then used GOrilla (*79*) to test for enrichments in the combined gene lists. Because gene function is related to evolutionary variance, for the background set we used the appropriate list of all high or low expression variance genes expressed at > 5 TPM.

#### Essential, haploinsufficient, and disease gene sets

The following gene lists were downloaded from the McArthur Lab gene lists repository at https://github.com/macarthur-lab/gene_lists: essential in culture, essential in mice, ClinGen haploinsufficient genes, Genes with any disease association reported in ClinVar, and neuromusclar disease genes.

Rare, single genes contributing to non-syndromic autism spectrum disorder were downloaded from the SFARI database at https://gene.sfari.org/ by selecting Category 1 genes (rare single gene variants, disruptions/mutations, and submicroscopic deletions/duplications directly linked to ASD) with a gene score of 1 (high confidence), 2 (strong candidate) or 3 (suggestive evidence).

Genes contributing to congenital heart disease were curated by filtering for genes annotated with “Congenital heart defects” in OMIM’s Morbid Map at https://omim.org/downloads/ as well as genes associated with congenital heart disease (DOID:1682) from the MGI Disease Ontology Browser at http://www.informatics.iax.org/disease.

### Samples for neuromuscular disease dataset

The cohort of neuromuscular disease patient RNA-seq described in this study is a superset of that described in Cummings et al. 2017 (*52*) (dbGaP accession phs000655.v3.p1) and 30 additional patients. Tissues were procured under Institutional Review Board (IRB) approved protocols at National Institue of Neurological Disorders and Stroke (Protocol #12-N-0095), Newcastle University (CF01.2011), Boston Children’s Hospital (03-12-205R), University College London (08ND17), UCLA (15-001919), and St. Jude Children’s Research Hospital (10/CHW/45). Patients were consented to these protocols in clinic visits prior to biopsy. Patient muscle biopsies were collected as described in Cummings et al. 2017 (*52*).

### RNA-seq for neuromuscular disease dataset patient data

RNA-seq from muscle biopsies was performed as described in Cummings et al. 2017 (*52*). Briefly, muscle biopsies or RNA were shipped frozen from clinical centers via a liquid nitrogen dry shipper and stored in liquid nitrogen cryogenic storage. All samples analyzed with H&E showed muscle quality sufficient to proceed to RNA-seq. RNA was extracted from muscle biopsies via the miRNeasy Mini Kit from Qiagen per kit instructions. All RNA samples were measured for quantity and quality and samples had to meet the minimum cutoff of 250 ng of RNA and RNA Quality Score (RQS) of 6 to proceed with library prep. RNA-Seq library preparation was performed at the Broad Institute Genomics Platform using the poly-A selection of mRNA with an Illumina TruSeq kit. Paired-end sequencing was performed in the Genomics Platform on Illumina HiSeq 2000 instruments. Read length and sequence coverage information is available in **table S4**.

GTEx BAM files were downloaded from dbGaP under accession ID phs000424.v6.p1 and realigned after conversion to FASTQ files with Picard SamToFastq. Both patient and GTEx reads were aligned using Star 2-Pass version v.2.4.2a using *hg19* as the genome reference and Gencode V19 annotations. Duplicate reads were marked with Picard MarkDuplicates (v.1.1099 available at http://broadinstitute.github.io/picard.

### Detecting outlier expression in patient samples

Genes expression values (log_10_(TPM)) were first normalized by TMM normalization (*76*) to the human skeletal muscle expression values used to originally to fit the OU parameters. For each gene in each patient sample, a z-score was calculated using the asymptotic mean and variance estimated from the evolutionary data. Z-scores were only calculated for genes that were assessed to fit better under the OU rather than the BM model (FDR < 0.05, see ***Fitting Ornstein-Uhlenbeck process parameters***) and whose asymptotic mean was estimated to be 5 TPM or higher. Z-scores were converted to p-values and then corrected for multiple hypothesis testing using the Benjamini-Hochberg FDR procedure (*78*). We used a FDR threshold of 0.01 to initially define significance. Of those, we removed another 330 genes that scored as a significant outlier in more than 25% of the GTEx samples.

As a comparator (**Fig. 3E, F**), z-scores were also calculated using the sample mean and variance estimated from health human GTEx samples. To ensure comparability between the two methods, we only calculated z-scores for genes that were not filtered out at any steps during the evolutionary method above.

### Detecting lineage-specific expression programs

Within each tissue, OU parameters for all four hypotheses (OU_all_, OU_primates_, OU_rodents_, OU_carnivores_) were estimated for each gene as described above. Models in which unrealistic parameters were estimated (optimal expression θ > 10^45^ or θ < 0) were removed. P-values were calculated using a likelihood ratio test comparing each of the OU models to the BM model. Results from each of the four hypotheses were then independently adjusted for multiple hypothesis testing using the Benjamini-Hochberg FDR procedure (*78*). For each gene, Akaike and Bayesian information criterion scores were calculated on all models that were significant against the null to determine the best fitting model. A gene was assigned to a best-fitting OU model only when both scores were in agreement. For additional stringency, we only defined genes as being differentially expressed if the fold change between the estimated means across the lineages (*e.g.*, primate mean vs. ancestral mean) was greater than 2-fold and if the mean reached at least 5 TPM in one of the lineages. To estimate the FDR, we performed the same procedure in each tissue using shuffled species assignments (on the same tree topology) and only retained hypotheses that achieved a FDR < 0.30. We performed GO enrichment analysis on each set of up- and down-regulated genes separately, using a background set of genes with mean expression of at least 1 TPM across all species in the appropriate tissue.

## Data availability

New RNA-seq data is available at GSE106077. Processed expression data, including expression values and estimated OU parameters for all species and tissues, is available at https://github.com/evee-tools.

## Acknowledgments

We thank Daniel MacArthur for help with patient data; Leslie Gaffney for artwork and advice on figures; and Hopi E. Hoekstra, Daniel L. Hartl, Yarden Katz, Akshay Krishnamurthy, Quinlan L. Sievers, Brian J. Haas, Atray Dixit, Rebecca H. Herbst, Dawn Thompson, Jessica Alföldi, and Carl G. de Boer for helpful discussions and feedback.

## Supplementary Figures

**Figure S1.**
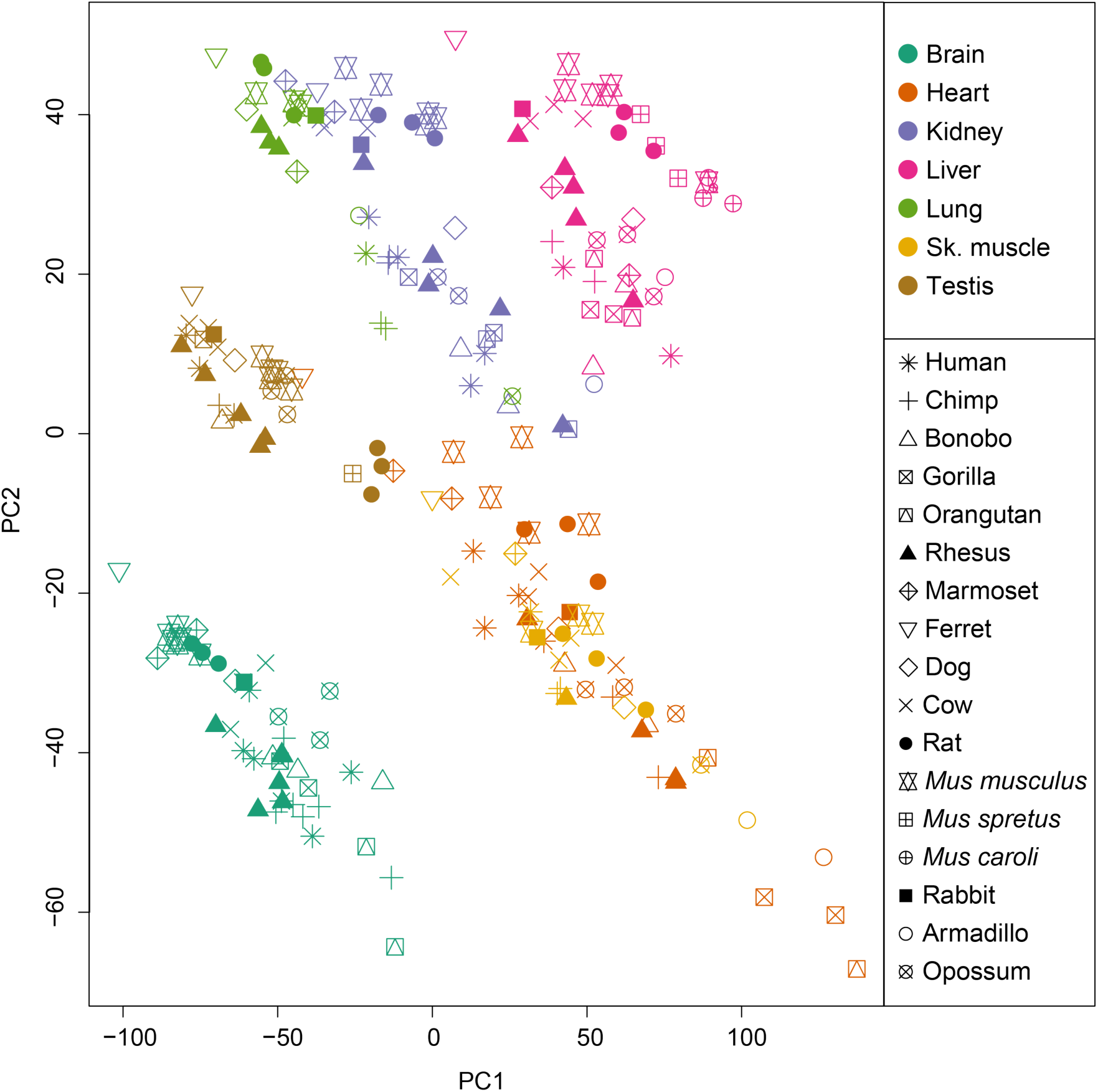
RNA-sequencing samples partition first by tissue, and then by species. Shown are the first two principal components (PCs) of a principal component analysis (PCA) of expression profiles of all 230 RNA-seq samples across 17 species (symbols) and 7 tissue types (colors).

**Figure S2.**
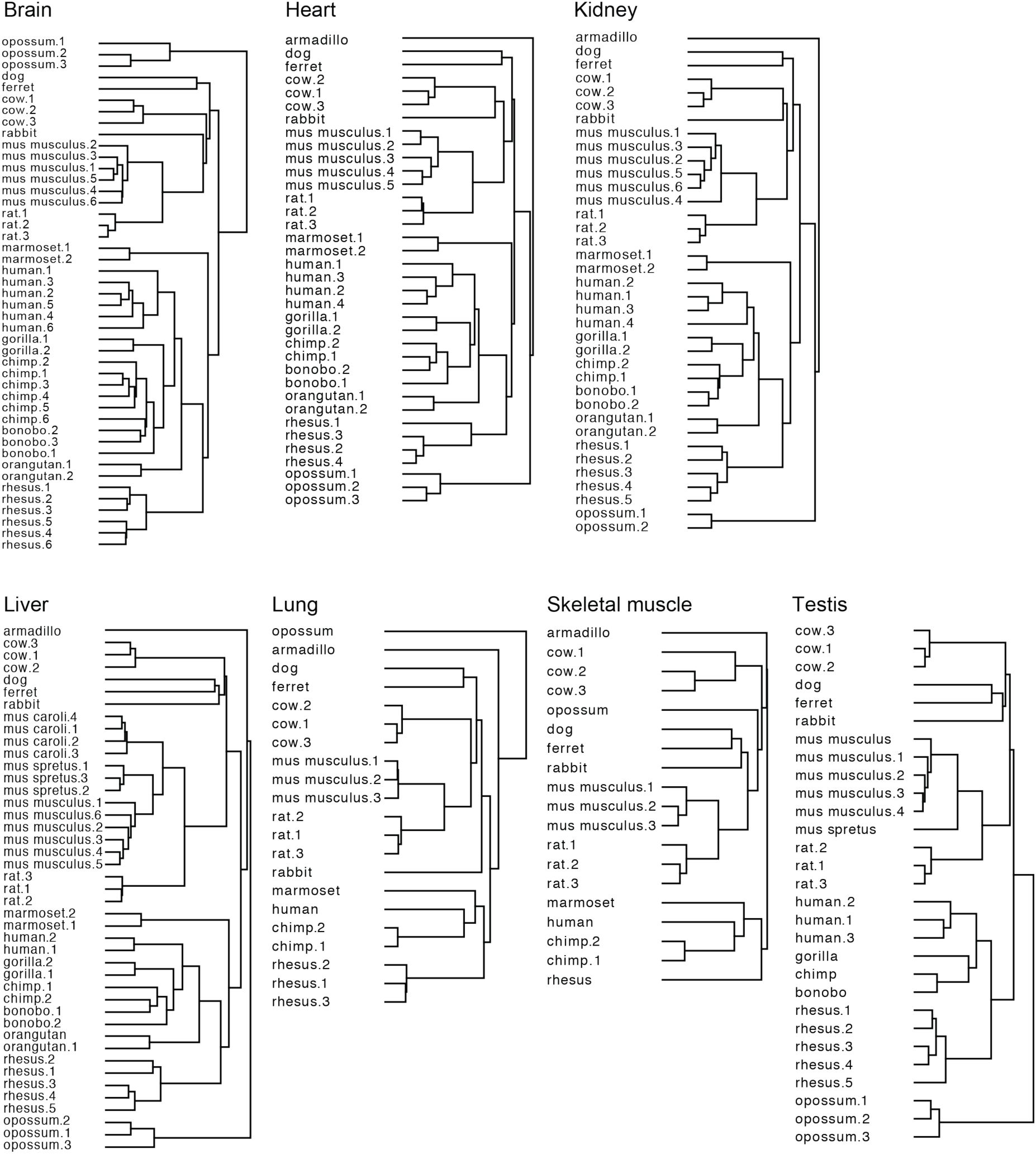
Hierarchical clustering of expression within tissues closely matches phylogenetic tree. Dendrograms from hierarchical clustering of gene expression (log_10_(TPM)) within each of seven tissue type (label, top) using Pearson’s correlation as the distance metric. See **Table S1** for detailed information about each sample.

**Figure S3.**
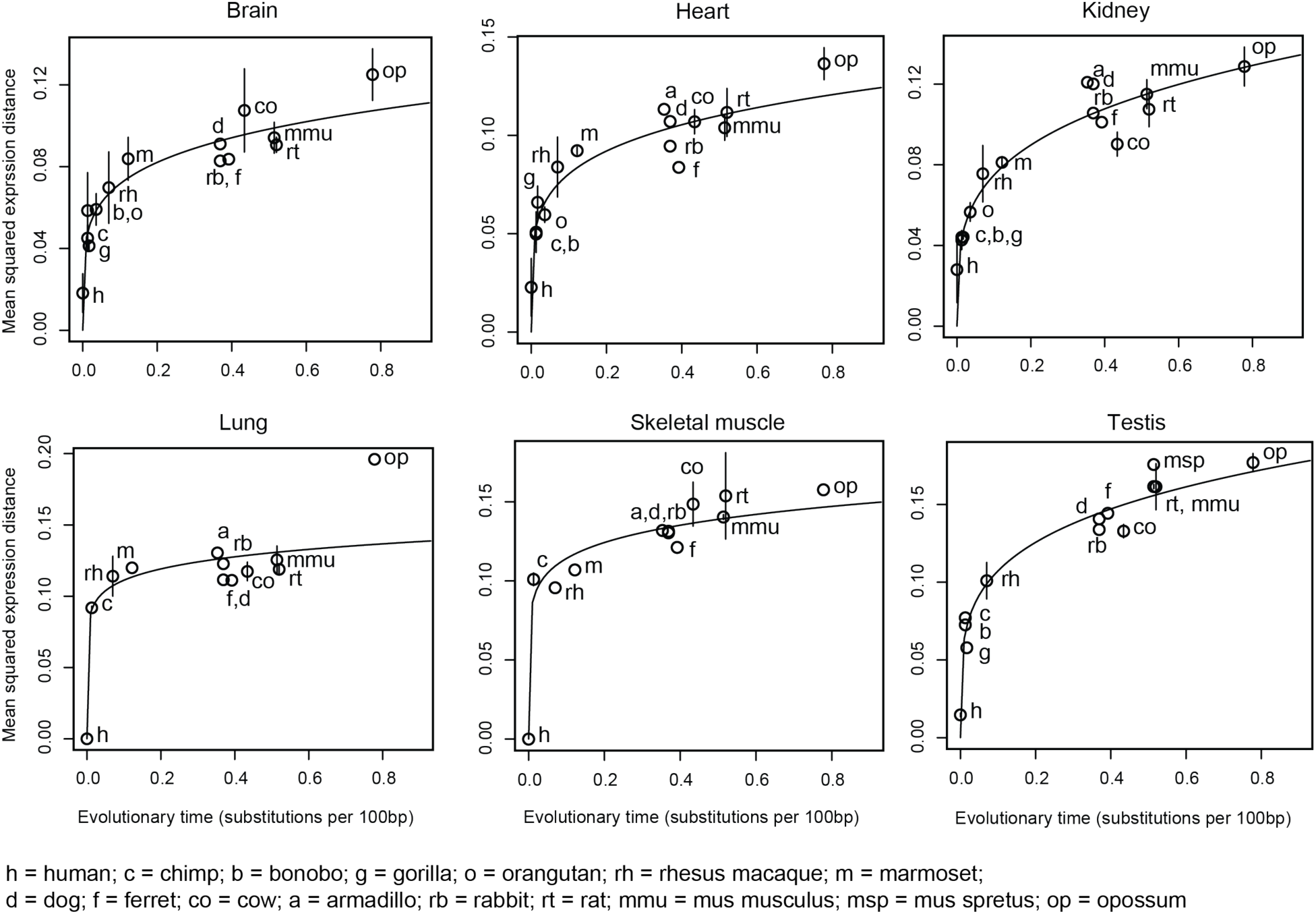
Expression evolution across mammalian lineages is accurately modelled by Ornstein-Uhlenbeck process. Shown are the pairwise mean squared expression distances (y-axis) between mammals and human for each of six tissue types (label, top) across evolutionary time, as estimated by substitutions per 100 bp (x-axis). Error bars: standard deviation of the mean across replicates. Solid line: Nonlinear regression fit. Bonobo, gorilla, and orangutan data is not available for lung and skeletal muscle.

**Figure S4.**
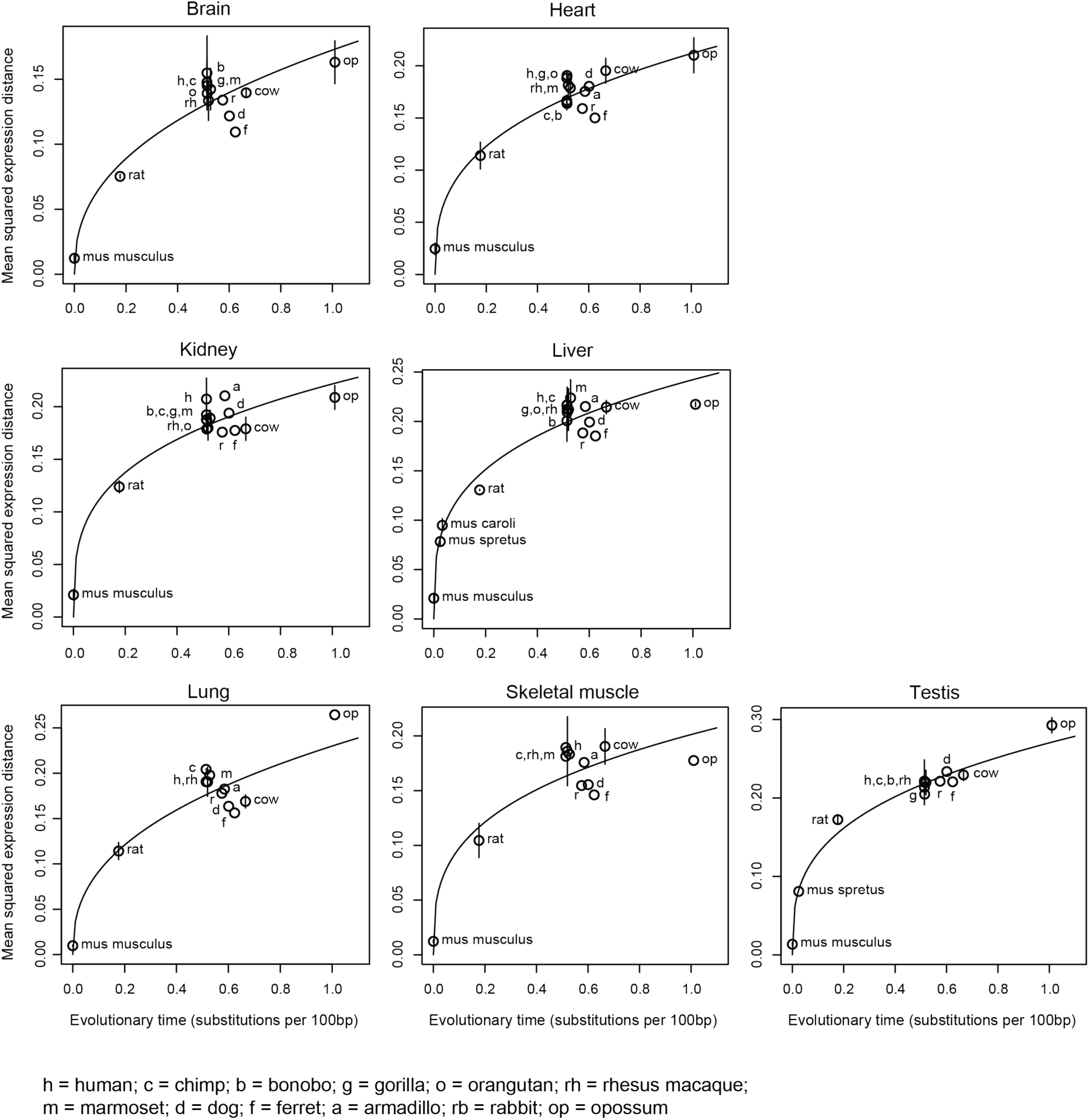
Expression evolution is accurately modelled by Ornstein-Uhlenbeck process, regardless of reference species. Shown are the pairwise mean squared expression distances (y-axis) between mammals and *mouse* for each of seven tissue types (label, top) across evolutionary time, as estimated by substitutions per 100 bp (x-axis). Error bars: standard deviation of the mean across replicates. Solid line: Nonlinear regression fit. Bonobo, gorilla, and orangutan data is not available for lung and skeletal muscle.

**Figure S5.**
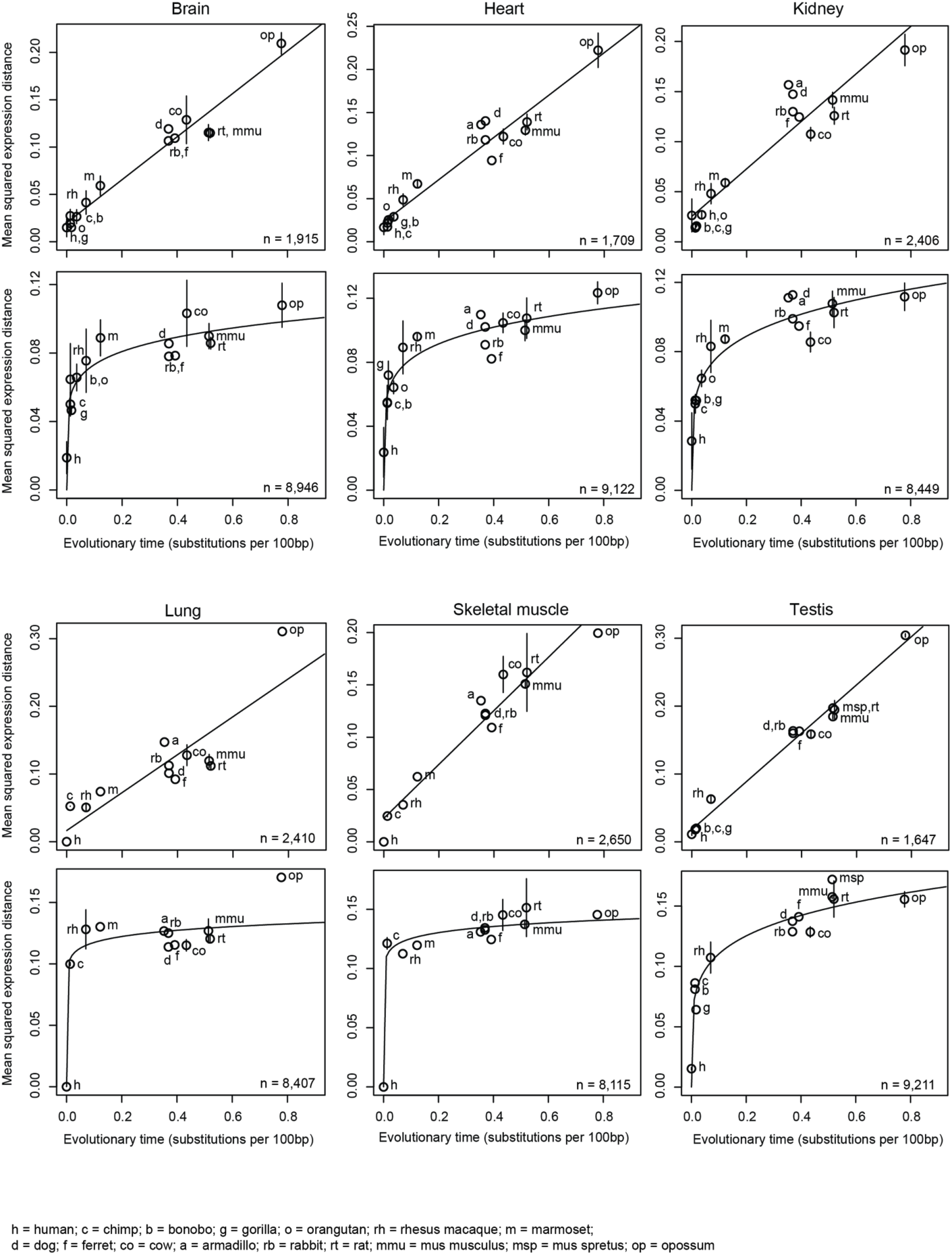
Modeling expression evolution by the Ornstein-Uhlenbeck process enables quantification of neutral and constrained selection on gene expression. Shown are the pairwise mean squared expression distances (y-axis) between mammals and human for each of six tissue types (label, top) across evolutionary time, as estimated by substitutions per 100bp (x-axis), separately for genes whose expression evolution fits better under a BM process (*i.e.* neutral evolution, top) and for genes whose expression evolution fits better an OU process (*i.e.* presence of stabilizing selection, bottom). Number of genes in each category is indicated on bottom right. Solid lines: Linear regression fit for BM genes and nonlinear regression fit for OU genes.

**Figure S6.**
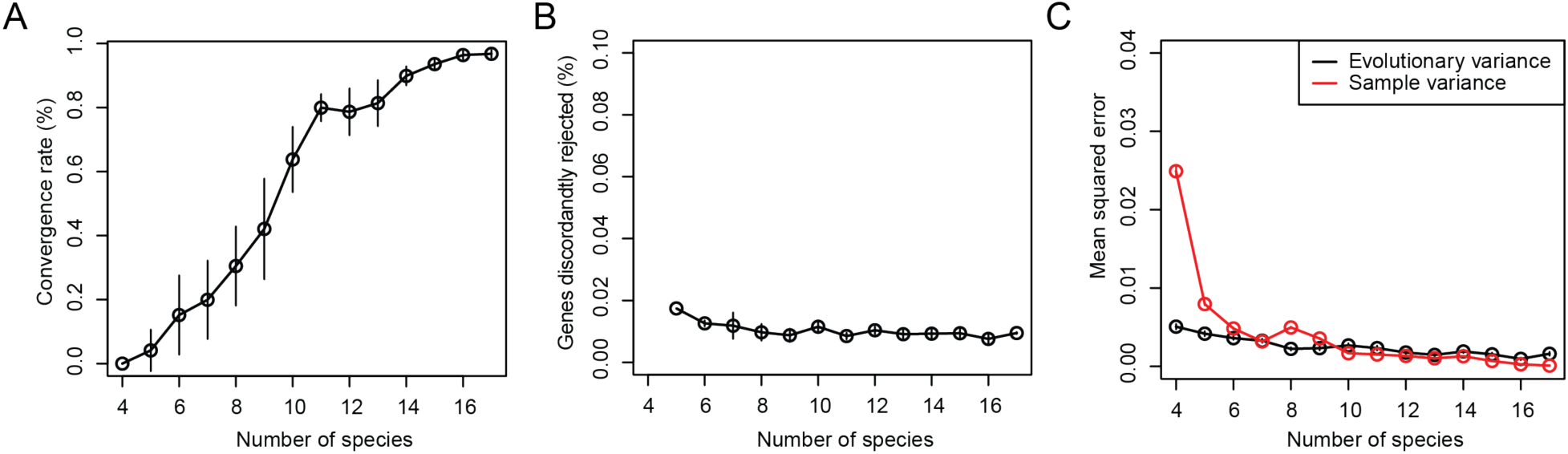
The Ornstein-Uhlenbeck process is a robust model for expression evolution. **(A)** Convergence with increased number of species. Percent of genes that reject the null hypothesis (BM process) (y-axis) when tested based on increasing number of species (x-axis) in the phylogeny, out of genes rejecting the null hypothesis when using the full phylogeny. Error bars: standard deviation of the mean across 10 iterations of sampled phylogenies. **(B)** Few genes are discordantly rejected. Percent of genes (y-axis) that reject the null hypothesis when using a sub-set of species (x-axis), but not when using all 17 species. Error bars: standard deviation of the mean across 10 iterations of sampled phylogenies. **(C)** Robustness of expression variance estimation. Mean squared error of variance estimated from OU model (black) or directly from the sample (red) based on increasing number of species (x-axis), compared to the variance was defined from results using the full phylogeny. Error bars: standard deviation of the mean across 10 iterations of sampled phylogenies.

**Figure S7.**
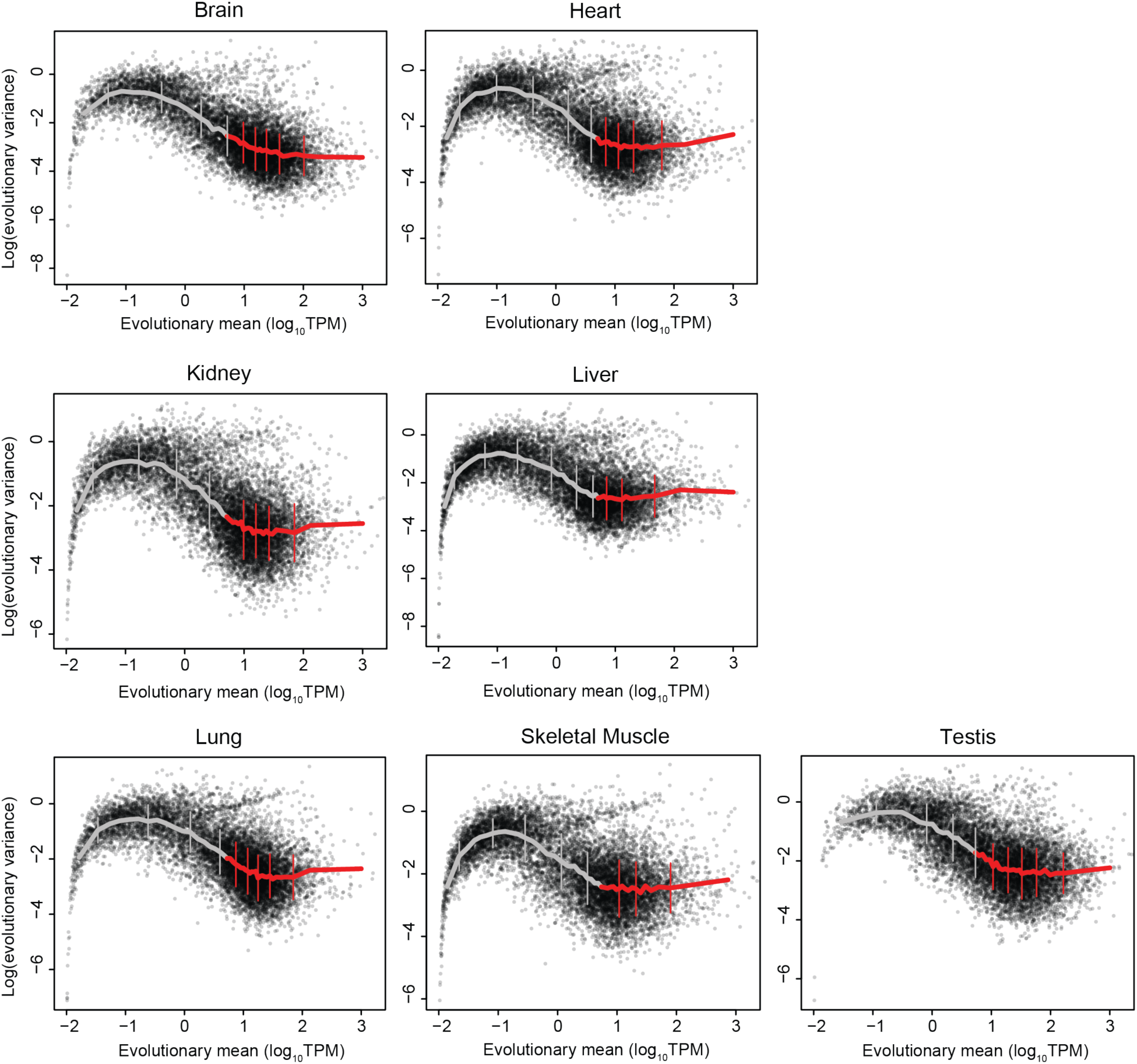
Lowly expressed genes have high estimates of evolutionary variance. Scatter plots show evolutionary mean (x-axis, log_10_TPM) and log(evolutionary variance) (y-axis) for each gene as estimated from OU model in each of 7 tissue types (label, top). Mean expression was binned into bins of 250 genes, and mean variance for each bin is shown in solid lines (grey/red: genes with expression lower/higher than 5 TPM). Error bars: standard deviation of the mean.

**Figure S8.**
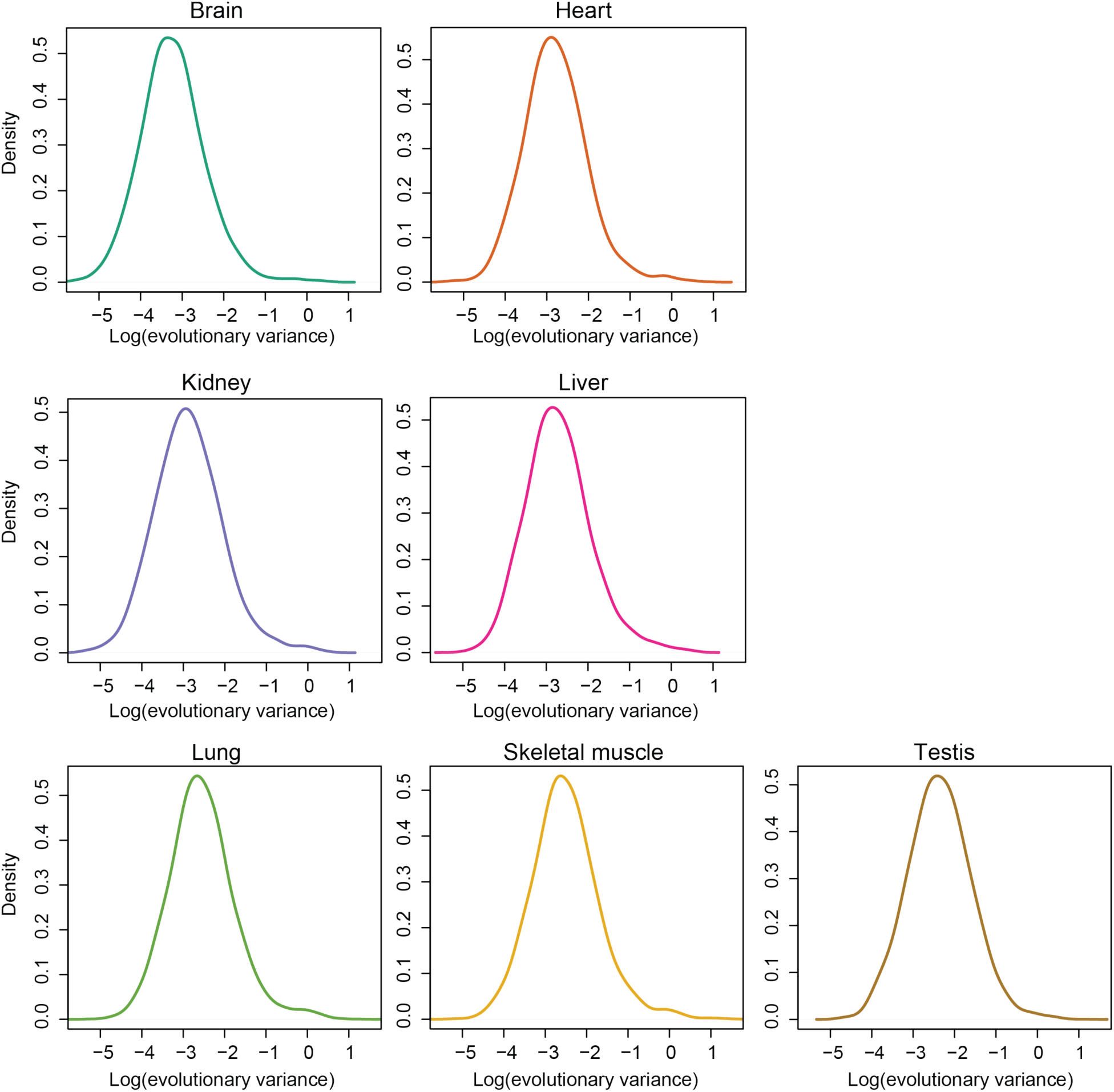
Evolutionary variance is log-normally distributed. Density plots of lo(evolutionary variance) for all genes in each of 7 tissue types (label, top).

**Figure S9.**
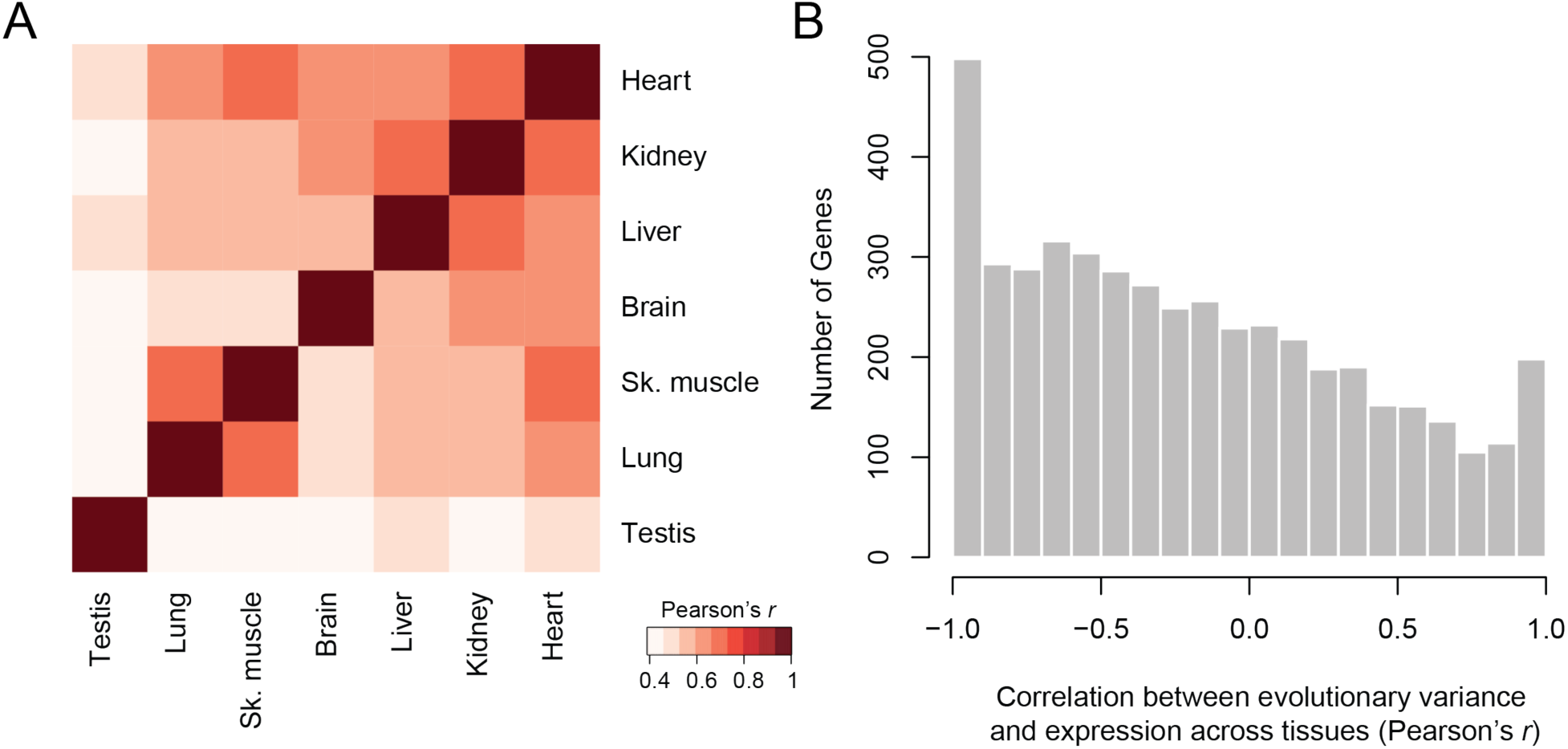
Evolutionary variance is correlated across tissues and negatively correlated with expression level. (**A)** Correlation of evolutionary variance across tissues. Heatmap of Pearson’s correlation coefficient of evolutionary variance estimates for all expressed genes (> 5 TPM) between each pair of tissues. (**B**) Negative correlation between evolutionary variance and expression level. Histogram shows the distribution of Pearson’s correlation coefficients between variance and expression across tissues, for genes expressed (> 5 TPM) in three or more tissues. Only variance and expression values for genes expressed at > 5 TPM were used for this analysis.

**Figure S10.**
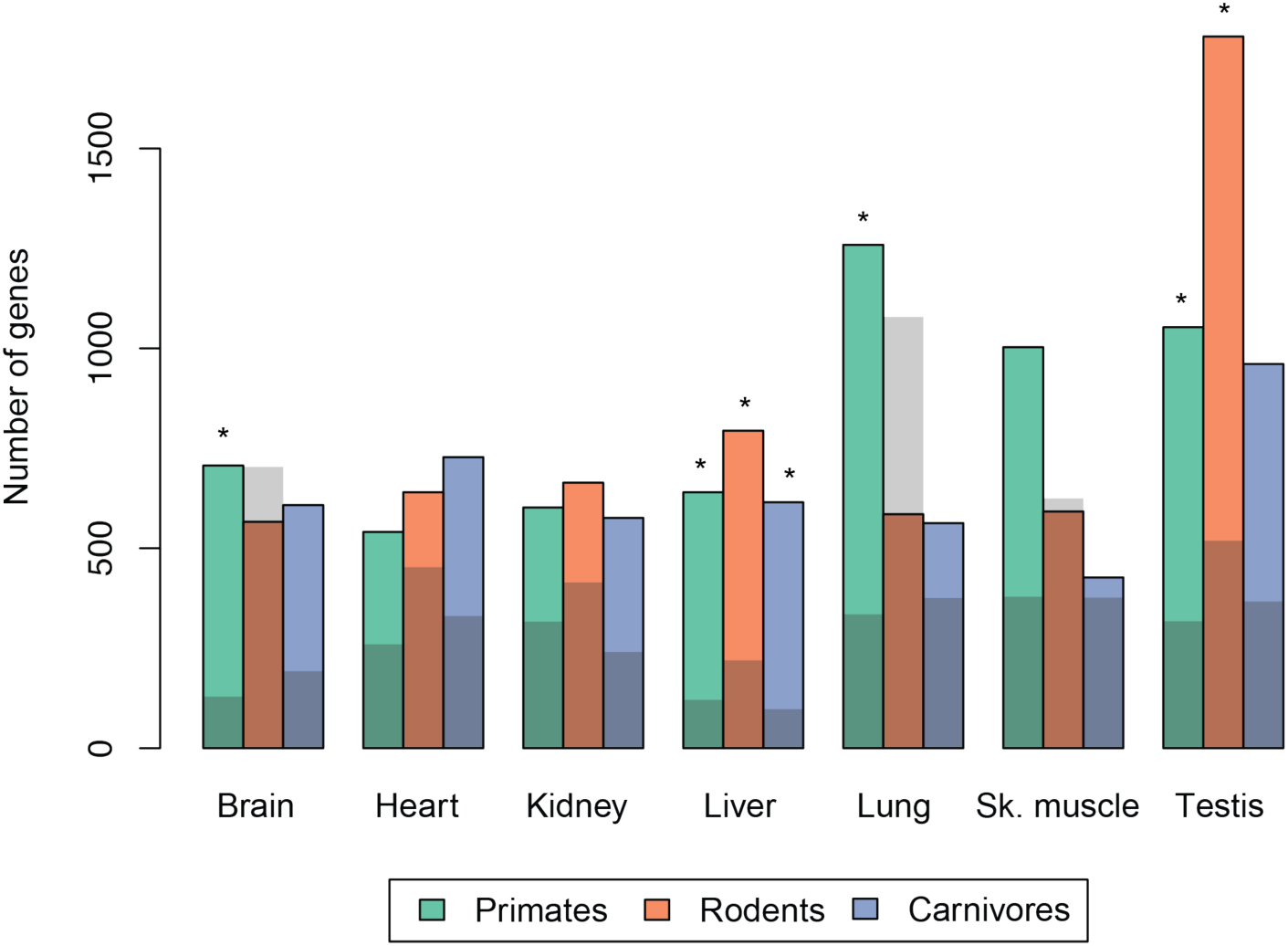
Permutation-based estimation of false discovery rates of genes with lineage-specific expression programs. Bar plot of number of genes (y-axis) detected to have lineage-specific expression changes across primates (green), rodents (orange), and carnivores (purple) in each of 7 tissue types (x-axis). Gray shading: number of genes detected by the same analysis when using a shuffled phylogeny (**SOM**). * denotes FDR < 0.30.

## Supplementary Tables

**Table S1. Data accessions and alignment statistics**

**Table S2. Gene ontology enrichments by evolutionary variance and tissue**

**Table S3. Gene ontology enrichments by evolutionary variance and sequence conservation**

**Table S4. RNA-seq statistics of muscle biopsies from patients with neuromuscular disease**

**Table S5. Gene ontology enrichments of genes with lineage-specific expression changes**

